# Elucidating pathogen interactions in *Tanacetum cinerariifolium* (pyrethrum) using fluorescently labelled *Didymella tanaceti* and *Stagonosporopsis tanaceti*

**DOI:** 10.64898/2026.03.30.715422

**Authors:** V. Carrillo, P.W.J. Taylor, A. Idnurm, T.L. Pearce, J.B. Scott, N. Vaghefi

## Abstract

Australia is the largest producer of Pyrethrum (*Tanacetum cinerariifolium*) globally. Amongst the constraints on production are the fungal pathogens *Didymella tanaceti* and *Stagonosporopsis tanaceti,* which pose a significant threat to the industry, causing substantial yield losses. While the infection biology of *S. tanaceti* is well characterised, knowledge of *D. tanaceti* and its potential interaction with *S. tanaceti* on plants remains limited, hindering disease management. We developed fluorescently labelled strains of both pathogens via *Agrobacterium tumefaciens*-mediated transformation (ATMT). Binary vectors carrying the mNeonGreen or tdTomato fluorescent protein genes were introduced into *D. tanaceti* and *S. tanaceti*, respectively, and expression of the fluorescent proteins was confirmed by microscopy. Genome sequencing revealed single-copy T-DNA insertions in all transformants, with minor genomic rearrangements at insertion sites. Detached leaf assays demonstrated that transformed strains retained pathogenicity, producing disease symptoms indistinguishable from those of the wild type. These fluorescently labelled variants enabled detailed visualisation of *D. tanaceti* infection biology and its interactions with *S. tanaceti*, including co-infection dynamics. Co-infection assays using fluorescent strains further facilitated simultaneous visualisation and differentiation of both pathogens within host tissues. Importantly, these tools also allowed the first description of the early stages of infection by *D. tanaceti* in pyrethrum leaves. This study represents the first successful transformation of *D. tanaceti* and *S. tanaceti*, providing valuable resources to investigate their infection processes.

## Introduction

Pyrethrum (*Tanacetum cinerariifolium*) is a perennial crop from the Asteraceae family, cultivated as the primary source of pyrethrins, which are natural insecticidal compounds (1). Pyrethrins are known for their ability to induce rapid insect knockdown, whilst having low toxicity in mammals, and rapid degradation in the environment (2). Australia is currently the largest global producer, contributing over 60% of the world supply, with most cultivation concentrated in Tasmania (3, 4).

During its 18-month cultivation cycle between sowing and first harvest, pyrethrum is susceptible to several fungal pathogens. Currently, *Didymella tanaceti* is the most prevalent foliar fungal pathogen (5), followed by *Stagonosporopsis tanaceti* (6), both causing significant yield losses (7, 8). *Didymella tanaceti* causes the disease tan spot and *S. tanaceti* causes ray blight (5, 9). These ascomycetes primarily cause leaf lesions. Historically, *S. tanaceti* was the dominant pathogen until 2009, and its biology and epidemiology are now well understood, enabling the development of targeted management strategies (9-12). However, a significant knowledge gap exists regarding the infection biology of *D. tanaceti*. Closing this gap is critical for developing specialised management strategies for tan spot.

Histological approaches have long been central to understanding plant-pathogen interactions, with staining techniques enabling the visualisation of pathogens *in planta* and the characterisation of infection structures. Traditional histopathology techniques provide key insights into how pathogen invade, colonise, and damage host tissues; however, they lack the specificity to rapidly distinguish individual species in mixed infections (13). This limitation is particularly relevant given that plants in natural environments are rarely infected by a single pathogen species (14). In contrast, fluorescently labelled pathogens provide a powerful means to track infection processes of specific pathogens *in planta* (13, 15, 16), enabling clear differentiation of the target pathogen from other microorganisms simultaneously colonising host tissues.

In nature, *Agrobacterium tumefaciens* is a plant-pathogenic bacterium that transfers a segment of DNA (T-DNA) from its Ti plasmid into the host plant genome, leading to the formation of crown galls (17). Advances in molecular techniques represented a critical tool to better understand the exact mechanism by which the DNA segment was transferred from the bacteria to the plant (18), leading to the first *Agrobacterium*-mediated transformation (ATMT) of the yeast *Saccharomyces cerevisiae* in 1995 (19, 20). Since 1998, ATMT has been extensively used in filamentous fungi to investigate gene function (21, 22) and to insert fluorescent proteins, such as DsRed and EGFP, for studying plant-pathogen interaction (15, 16, 23, 24). The green fluorescent protein (GFP), derived from *Aequorea victoria* jellyfish, is a highly stable reporter capable of resisting temperatures over 60 °C, low pH and proteases (23, 25). However, this wild type GFP is poorly translated in fungi, leading to the development of codon-optimised variants such as SGFP and EGFP, with SGFP preferred for filamentous fungi (23). Over 10 years ago, the discovery of the yellow fluorescent protein LanYFP from the lancelet *Branchiostoma lanceolatum* led to the engineering of mNeonGreen, a monomeric, bright green fluorescent protein (26) that has been expressed in filamentous fungi such as *Aspergillus fumigatus* (27) and *Sordaria macrospora* (28). Similarly, the discovery of red fluorescent proteins expanded the available spectral range for live imaging. The original red fluorescent protein, DsRed from *Discosoma* spp., was later engineered into monomeric and dimeric variants with enhanced brightness and photostability, such as mRFP1 and tdTomato (29, 30). The latter has been successfully incorporated into *Sclerotium rolfsii* (31) and *Aspergillus niger* (32), producing a characteristic bright red-yellow fluorescence.

This study aimed to transform *D. tanaceti* and *S. tanaceti* via ATMT to express mNeonGreen and tdTomato, respectively. The infection capacity and transformant stability of selected transformed strains were also corroborated. These fluorescently labelled variants enabled the detailed visualisation of *D. tanaceti* infection biology and its interactions with *S. tanaceti*, including co-infection dynamics. Hence, this report provides a first look into the early stages of the infection process of *D. tanaceti* and demonstrates the utility of fluorescently labelled pathogens as a powerful tool to distinguish between two pathogens within the same leaf tissue.

## Materials and methods

### Fungal cultures and spore suspension preparation

For this study, strain BRIP 61988 (33) was selected as the representative wild type of *D. tanaceti*, and isolate UOM ST2 were selected as the representative isolate of *S. tanaceti*. Pure cultures of each species were obtained by single sporing. Shortly, upon sporulation, 5 ml of autoclaved reverse osmosis (RO) water was added to each plate, and the surface was scraped with a sterile scalpel blade. The resulting spore suspensions were filtered using an autoclaved funnel and a single layer of Miracloth® (Millipore, Germany). Spore suspensions of each species were adjusted to 1 × 10^5^ spores/ml with a hemocytometer (Livingston, Australia) and spread across a Petri plate containing water agar (WA) (Supplementary material). In the dissecting microscope (Leica Microsystems, Germany), single spores were picked up and transferred to different media. *Didymella tanaceti* pure cultures were grown on potato dextrose agar (PDA, Difco, Australia), and *S. tanaceti* pure cultures were grown on V8 juice agar (V8) (12) (Supplementary material). All cultures were kept at 21°C at a 12-hour light and dark cycle for 20 days.

### *Agrobacterium*-mediated transformation

#### Plasmid mapping

Targeted isolates of *D. tanaceti* and *S. tanaceti* were transformed using the binary vectors pMAI31 (111,27 bp) and pMAI32 (118,47 bp) respectively. Both plasmids were constructed by modifying the T-DNA section of pLAU2 (34, 35) that contains the kanamycin resistance gene *kanR*. Both plasmids carried fluorescent protein genes: pMAI31 (Figure S1. A) contains *mNeonGreen* and pMAI32 (Figure S1. B) contains *tdTomato*, each driven by the *act1* promoter and *trp3* terminator from *Leptosphaeria maculans*. The hygromycin resistance gene *hygR* was driven by *trpC* promoter and terminator from *Aspergillus nidulans*.

#### Co-culture of Agrobacterium and fungi, and initial antibiotic selection

For the fungal transformation, spore suspensions from *D. tanaceti* and *S. tanaceti* wild type strains were obtained by adding 5 mL of autoclaved RO water to each plate. The surface was scraped with a sterile scalpel blade, and the resulting spore suspensions were filtered using an autoclaved funnel and a single layer of Miracloth®. The spore suspension obtained from each strain was adjusted to a concentration of 2-3 × 10^7^ spores/ml.

Liquid cultures of *Agrobacterium* containing the plasmids were diluted with LB to approximately optical density (OD) 600nm of 0.6. In a 2 ml tube, 400 μl of diluted *Agrobacterium* was mixed with 400 μl of fungal spore suspension. From the admixture, 400 μl was transferred to a 15 cm Petri plate containing induction media (IM) supplemented with 10 mM of acetosyringone (AS). Plates were incubated at 22°C in darkness for 2-3 days without sealing (Supplementary material). *Agrobacterium tumefaciens* harbouring plasmid pMAI31 was co-cultured with spore suspension of strain BRIP 61988 (*D. tanaceti*) and *A. tumefaciens* containing pMAI32 was co-cultured with the spore suspension of strain UOM ST2 (*S. tanaceti*).

Subsequently, in a biosafety cabinet, 25 ml overlay media (PDA for *D. tanaceti* and V8 for *S. tanaceti*) with 100 µg/ml cefotaxime (Fluorochem, United Kingdom) and 50 µg/ml hygromycin (Thermo Fisher Scientific, USA) was poured as an overlay and left to dry for 10 minutes. Plates were then sealed with parafilm and incubated at 22 °C under light. Colony formation was monitored daily for 10 days. Colonies were transferred onto fresh PDA or V8 plates containing hygromycin and cefotaxime for an additional selection round, placing a single colony per plate. After the cultures covered the 90 mm Petri plates, transformed strains were subcultured to antibiotic-free media.

#### Establishment of pure cultures and stability of transformants

*Didymella tanaceti* transformants were single-spored to establish pure cultures of the transformed strains. Spore suspensions of *D. tanaceti* transformants were prepared from 3-week-old transformed plates, adjusted to 1×10^5^ spores/ml by counting the spores on a hemocytometer and diluting the suspension with RO water. Then, 100 μl of spore suspension was placed on a Petri plate containing WA, and the content was spread using a sterile rod. After 24 hours, germinating spores were observed under the dissecting microscope, picked up with a sterile needle, transferred onto PDA and kept at 22°C at a 12-hour light and dark cycle. *S. tanaceti* transformants were hyphae-tipped by sampling single hyphae from the margin of one-week-old cultures and transferring to V8 agar. Cultures were kept at 22°C at a 12-hour light and dark cycle.

Two consecutive rounds of culturing on PDA and V8 agar media (for *D. tanaceti* and *S. tanaceti* transformant, respectively) containing hygromycin and cefotaxime were performed before subculturing in antibiotic-free media.

### Fluorescence microscopy and quantification

The fluorescence of spores and hyphae samples, obtained from pure cultures of transformed isolates, was observed by mounting a 5 μl spore suspension droplet and 2-3 mm^2^ agar plug on a glass slide. Fluorescence was confirmed using compound microscope Leica DM6000 (Leica Microsystems, Germany). The Yellow Fluorescent Protein filter (YFP, excitation: 514 nm, emission: 527 nm) was used to detect mNeonGreen and the DsRed fluorescent filter (excitation: 558 nm, emission: 583 nm) was used to detect tdTomato.

Hyphae fluorescence intensity, obtained from pure cultures of transformed isolates, was quantified using Fiji image processing software version 2.16.0/1.54p (36), and splitting the channels to measure the fluorescence in grey-scale values. Fifteen regions of interest (ROI) were measured per transformant by placing the ROI (w=1.96, h=1.96) in hyphae sections that did not contain any fluorescent hyphae in the background. The background noise was subtracted to obtain the fluorescence intensity values, which were then used to calculate the corrected total cell fluorescence (CTCF) value (37, 38).

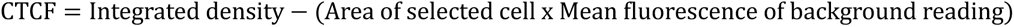

Images were captured at a 40× magnification with 1 s exposure. One-way ANOVA and Tukey post hoc test were used to compare fluorescence between samples. Based on CTCF values, *D. tanaceti* transformant UOM DT1 and UOM DT5, and *S. tanaceti* transformant UOM STC1R2 and UOM STC4R2, were selected for molecular characterisation.

### Growth rate and morphological characterisation

Growth rates were measured in triplicate on PDA (*D. tanaceti*) and V8 agar (*S. tanaceti*). A single agar plug was placed at the centre of a marked 90 mm Petri plate. All cultures were kept at 22°C at 12-hour light/dark cycle. Colony diameter was measured every three days along the same line until colonies covered the plate.

Spore shape and size were assessed by measuring the length and width of 30 spores from each of the five *D. tanaceti* transformants. Spore suspensions were prepared from two-week-old cultures kept at 22°C at 12-hour light/dark cycle, by placing 5 ml of autoclaved RO water and scraping the agar surface with a sterile blade. The liquid was then filtered using an autoclaved funnel wrapped with Miracloth into 50-ml centrifuge tubes. Spore suspensions were mounted on glass slides and examined under a compound microscope (Leica DM6000) at 100× magnification. Measurements were taken using the analysis tools available at the Leica Application Suite (LAS) software.

### Molecular characterisation of *S. tanaceti* and *D. tanaceti* transformants

Two transformed strains from each species were selected for whole-genome sequencing to determine the copy number of T-DNA insertions and their locations in the genomes. For *D. tanaceti*, transformed strains UOM DT1 and UOM DT5, and BRIP 61988 wild type were selected; for *S. tanaceti*, strains UOM STC1R2 and UOM ST4R2 were chosen for genome sequencing.

Subcultures were grown on potato dextrose broth (PDB, Difco, Australia) in a shaker at 25°C for two weeks. In the biosafety cabinet, mycelium samples were collected with sterilised tweezers and placed in an autoclaved paper towel (autoclave cycle: 121°C for 20 min) to remove the excess liquid. The mycelium was transferred to 2 ml safe-lock tubes and placed immediately in a cooler containing liquid nitrogen. Samples were freeze-dried for 48 hours in a freeze dryer (Biobase). Two magnetic beads were then added to each tube, and samples were lysed for 30 s at 30 Hz frequency in the TissueLyser III (Qiagen). DNA was extracted following the manufacturer’s instructions for the DNeasy Plant Mini Kit (Qiagen, Germany). DNA quality was assessed via gel electrophoresis (1.5% agarose), and concentration was measured using the Qubit 2.0 fluorometer (ThermoFisher Scientific).

Whole genome sequencing of the selected strains was performed using Illumina HiSeq platform at the Australian Genome Research Facility (AGRF, Melbourne) as paired-end reads of 150 bp. The NovaSeq Control Software (NCS) v1.3.0.39308 and Real-Time Analysis (RTA) v4.29.2 performed image analysis in real-time. Illumina DRAGEN BCL Convert 07.031.732.4.3.6 pipeline was used to generate the sequence data. Adapter sequences were removed with Trimmomatic (Galaxy Version 0.36.6) and the genome was assembled using SPAdes (Galaxy Version 4.1.0+galaxy0) (39). Assembly quality and completeness were assessed using Quast (Galaxy version 5.3.0+galaxy0) and BUSCO (Galaxy version 5.8.0+galaxy1) using the BUSCO v5 lineage datasets (odb10). The Kmer counting software Jellyfish v.2.3.1 (40) was used to estimate the genome size using raw Illumina data.

The number of T-DNA inserts of each strain was determined by Custom BLAST in Geneious Prime v2025.0.3 (https://www.geneious.com), using the T-DNA sequence as query against the assembled genomes of the transformed strains. Genes were predicted using AUGUSTUS (39) with *Botrytis cinerea* as the reference species. The predicted gene models were subjected to BLASTx searches against protein databases to infer putative gene functions. Insertion sites were identified by mapping the complete T-DNA against the 150 bp paired raw reads of each strain and BLAST analysis using the flanking regions as query against the annotated wild type genomes. Gene insertion sites were illustrated schematically using BioRender (https://biorender.com) to visualise the T-DNA integration and associated genomic features

### Detached leaves assay - *D. tanaceti* transformed strains analysis

A total of 46 pyrethrum leaflets was obtained from 7-month-old pyrethrum plants of the BR3 variety, grown in controlled temperature rooms set at 20°C, with 10 hours of light and 14 hours of darkness, at 60% percent humidity. The leaves were inoculated with spore suspensions of *D. tanaceti* wild type strain BRIP 61988 or the transformed strain UOM DT5. Spore suspensions were adjusted to 1×10^7^ spores/ml and supplemented with Tween 20 (0.02% v/v). For inoculation, a 5-10 µl droplet was placed at the centre of each leaflet and spread across the surface with the pipette tip. Control leaves received RO water containing Tween 20 (0.02% v/v). Following inoculation, samples were placed in glass Petri plates containing moistened sterile paper towels to maintain humidity and incubated at 22°C under a 12 h light/12 h dark cycle in a growth cabinet.

Five leaflets per strain were collected at 24, 48, 72 hours after inoculation (HAI) and at 5 days after inoculation (DAI) to examine germination and early infection. At 24 and 48 HAI, samples were observed from the adaxial surface using a compound microscope (Leica DM 6000). Leaflets collected at 72 HAI and 5 DAI were transversely sectioned by hand with a razor blade, with infected tissues stabilised between polystyrene sheets during sectioning.

For the samples inoculated with the wild type, tissues were cleared in a 1:1 (v/v) ethanol-glacial acetic acid solution for 12 hours and stained with lactophenol cotton blue to visualise fungal structures before observation under brightfield microscopy. Samples infected with the fluorescent strain UOM DT5 were examined directly under the compound microscope using the YFP filter set.

### Detached leaves assay - *S. tanaceti* transformed strains analysis

The bioassay required ten 7-month-old leaves collected from pyrethrum plants (BR3 variety), grown in controlled temperature rooms set at 20°C, with 10 hours of light and 14 hours of darkness, at 60% percent humidity. Leaves were inoculated with the wild type strain UOM ST2 or transformed strains STC1R2 of *S. tanaceti*. A 10 μl droplet of spore suspension adjusted to 1×10^7^ spores/ml, was placed at the centre of each leaflet and spread across the surface with a pipette tip. Samples were placed on a glass Petri plates containing moistened sterile paper towels to maintain humidity at 22°C at 12-hour of light and dark cycle in a growth cabinet.

Leaflets were sampled 3 DAI and hand-sectioned transversally using a dissecting microscope. Sectioned samples were observed under the compound microscope equipped with DsRed fluorescent filter to visualise infected tissue and the pathogen *in planta*. *S*. *tanaceti* transformant were considered phenotypically identical to the wild type if initial symptoms appeared at the same time and fluorescent mycelia were observed in the inner lamella at 3 DAI (41).

### Detached leaves assay - *D. tanaceti* and *S. tanaceti* co-infection

Ten 7-month-old leaflets were co-inoculated with the transformed strain STC1R2 of *S. tanaceti* and the transformed strain UOM DT5 of *D. tanaceti*. At 7 DAI, leaves were sampled and transversely sectioned using a razor blade, with infected tissues stabilised between polystyrene sheets during cutting. Sections were mounted in droplets of RO water on glass slides and examined under a compound microscope equipped with DsRed and YFP fluorescence filter sets. Images of infected plant tissue were processed using Fiji.

## Results

### Agrobacterium-mediated transformation of D. tanaceti and S. tanaceti

From the co-culture of *Agrobacterium* harbouring the plasmids and the spore suspensions of the wild type isolated BRIP 61988 and UOM ST2, five hygromycin-resistant colonies of *D. tanaceti* and four of *S. tanaceti* were recovered. The transformant strains of *D. tanaceti* were designated as UOM DT1, UOM DT2, UOM DT3, UOM DT4 and UOM DT5, and the four transformant strains of *S. tanaceti* were designated as UOM STC1R2, UOM STC2R2, UOM STC3R2, and UOM STC4R2. All transformant strains of both species consistently retained fluorescence following subculturing, indicating successful and stable integration of the construct.

### Fluorescence microscopy and CTCF values

Fluorescence in both species was confirmed under the compound microscope, using the YFP filter to observe *D. tanaceti* strains containing the mNeonGreen fluorescent protein, and DsRed filter to observe the *S. tanaceti* strains containing tdTomato fluorescent protein. Spores and hyphae of both species exhibited strong fluorescence. *Didymella tanaceti* transformant strains expressing mNeonGreen displayed green fluorescence (Figure 1. A), with variable intensities among strains, while the *S. tanaceti* transformant strains expressing tdTomato exhibited red-orange fluorescence (Figure 1. B) with no apparent difference between strains. Fluorescence was strain-specific, with no signal detected outside the corresponding channels. *Didymella tanaceti* transformants expressing mNeonGreen did not emit fluorescence in the DsRed channel, while *S. tanaceti* transformants expressing tdTomato showed no fluorescence in the green channel.

**Figure 1.**
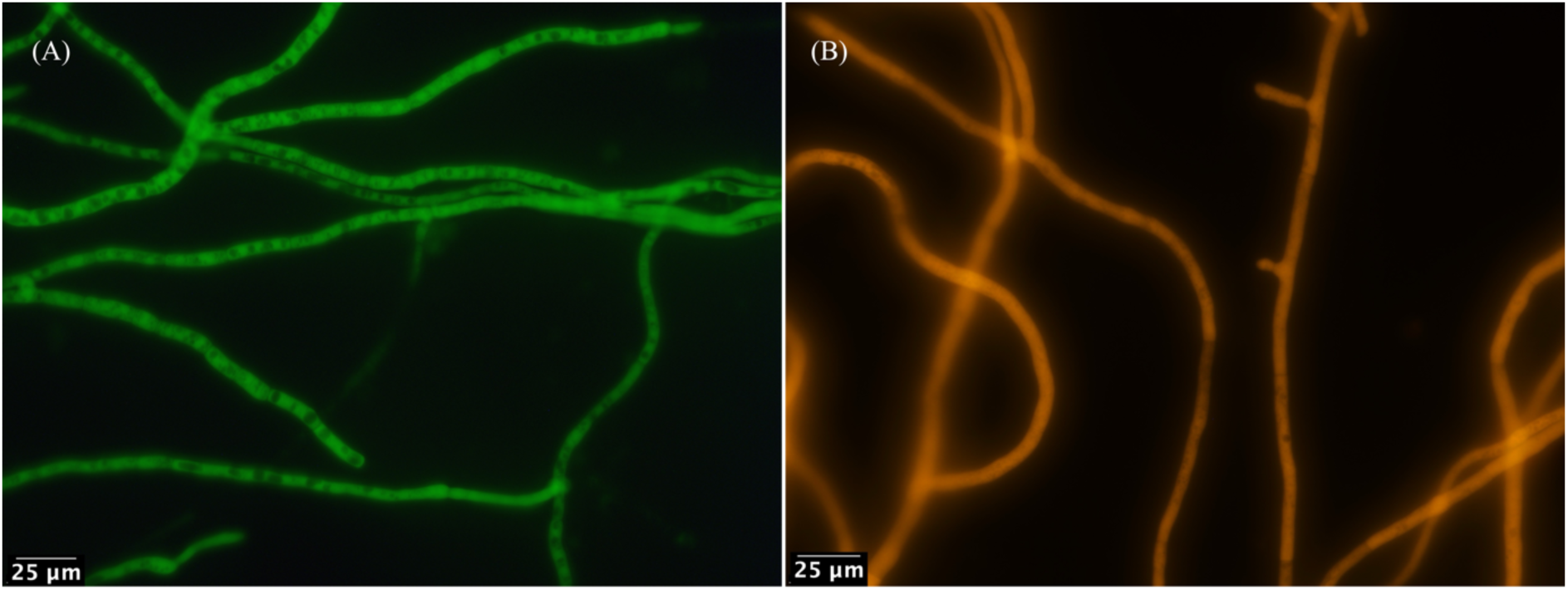
Fluorescent microscopy images of transformed *Didymella tanaceti* and *Stagonosporopsis tanaceti*. (A) Mycelium of strain UOM DT5 observed under a YFP filter (excitation 513 nm, emission 527 nm) showing expression of the mNeonGreen fluorescent protein. (B) Mycelium of strain UOM STC1R2 observed under a DsRed filter (excitation 558 nm, emission 583 nm) showing expression of the tdTomato fluorescent protein. Scale bar = 25 μm Microscopically, no apparent differences in fluorescence intensity were observed among the four *S. tanaceti* transformant strains. In contrast, slight variations were noted among *D. tanaceti* transformant strains. These differences were quantified using CTCF values derived from greyscale measurements. Statistical analysis revealed no significant differences among *S. tanaceti* transformant strains (Figure 2. A). In *D. tanaceti*, however, strain DT5 exhibited a significantly higher CTCF value (p = 0.02 - 0.0002) compared with the other three transformant strains, indicating greater fluorescence intensity (Figure 2. B).

**Figure 2.**
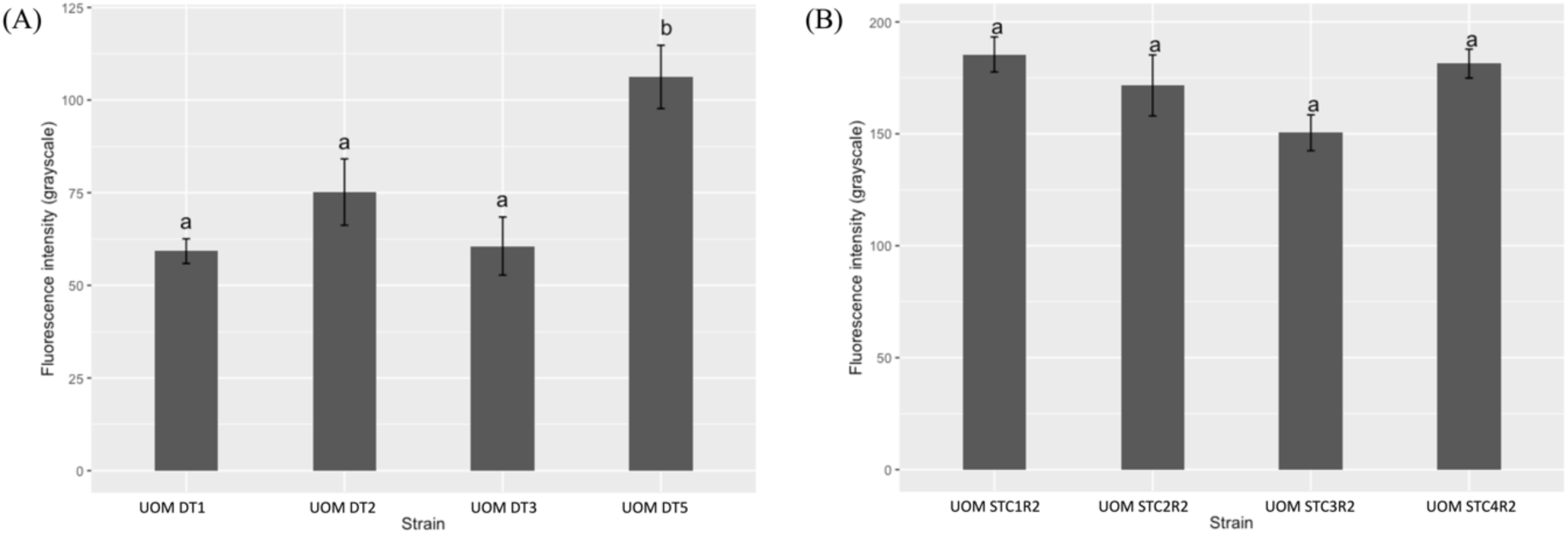
Fluorescence intensity of transformed strains of *Didymella tanaceti* and *Stagonosporopsis tanaceti* based on greyscale-derived corrected total cell fluorescence (CTCF) values. Different letters indicate statistically significant differences (p < 0.05). (A) *D. tanaceti*: strain UOM DT5 exhibited significantly higher fluorescence intensity than the other strains (p = 0.0002). (B) *S. tanaceti*: no significant differences were observed among transformed strains (p > 0.05).

### Morphological comparison against the wild type

Colony morphology remained the same across cultures. *D. tanaceti* wild type and transformed strains exhibited the characteristic green olivaceous mycelia (Figure S2. A-F), with sporulation observed after seven days. Likewise, *S. tanaceti* wild type and transformant developed white mycelia during the first week (Figure S3. A-E), with pycnidia appearing three weeks post-subculturing. Colony growth rates for *S. tanaceti* were consistent across all strains, with complete coverage of 90 mm Petri plates achieved within nine days (Table S1). In contrast, one *D. tanaceti* transformed strain (UOM DT4) displayed significantly reduced growth rate compared to the wild type (p = 0.008) and was, therefore, excluded from subsequent analyses (Table S2).

Spore shape of *D. tanaceti* transformed isolates was cylindrical to ellipsoidal with rounded ends, rarely septate. For UOM DT1, spores measured ranged 4.3–7.1 × 2.6–4.2 µm; UOM DT2, 5.3–7.6 × 2.7–4.7 µm; UOM DT3, 4.3–7.3 × 2.5–4.0 µm; UOM DT4, 5.0–8.0 × 1.3–4.7 µm; and UOM DT5, 5.9–8.5 × 2.9–4.7 µm

### Molecular characterisation T-DNA and insertion sites

The Assembly size of the wild type *D. tanaceti* strain BRIP 61988 was 41.1 Mbp, with a sequencing coverage of 141×, considering contigs >1,000 bp in length. The transformed strains UOM DT1 and UOM DT5 had an assembly size of 39.4 Mbp and 38.9 Mbp, with sequencing coverages of 205× and 160×, respectively. The estimated genome size for *D. tanaceti* isolates was 40.7 Mbp using Jellyfish. For *S. tanaceti* transformed strains STC1R2 and STC4R2, the assembly sizes were 33.3 Mbp and 33.4 Mbp, respectively, considering contigs >1,000 bp in length, and the sequencing coverages were 68.6× and 73.5×, respectively. The estimated genome size for *S. tanaceti* was 35.5 Mbp.

BUSCO analysis against 6,641 orthologous groups indicated high completeness for the *D. tanaceti* wild type and transformed strains. The wild type strain BRIP 61988 recovered 95.5% of expected genes (6,341 complete BUSCOs), of which 95.2% (6,325) were single-copy and 0.2% (16) were duplicated, while 1.3% (84) were fragmented and 3.3% (216) were missing. A similar completeness was observed in the transformed strains UOM DT1 and UOM DT5. The former, displayed 95.5% completeness (6,341 complete BUSCOs), with 95.2% (6,325) single-copy, 0.2% (16) duplicated, 1.3% (84) fragmented, and 3.3% (216) missing. The latter showed a nearly identical profile, with 95.6% of genes detected (6,346 complete BUSCOs), comprising 95.3% (6,330) single-copy and 0.2% (16) duplicated, alongside 1.2% (78) fragmented and 3.3% (217) missing.

For *S. tanaceti* transformed strains, BUSCO analysis against 6,641 orthologous groups indicated high completeness. Strain UOM STC1R2, had 96.1% of expected genes detected (6,381 complete BUSCOs). Of these, 95.9% (6,370) were present as single-copy and 0.2% (11) as duplicated, while 1.2% (78) were fragmented and 2.7% (182) were missing. Similarly, for strain UOM STC4R2, 96.0% of genes recovered (6,374 complete BUSCOs), comprising 95.8% (6,364) single-copy and 0.2% (10) duplicated, with 1.2% (80) fragmented and 2.8% (187) missing.

The T-DNA construct in *D. tanaceti* transformants carrying the mNeonGreen fluorescent protein gene was 4,951 bp in length, whereas the T-DNA construct in *S. tanaceti* transformant carrying the tdTomato fluorescent protein gene was 5,671 bp. BLASTn searches of the T-DNA sequence against each transformed genome assembly identified contigs containing the T-DNA. Strains UOM DT1, UOM STC1R2, and UOM STC2R2 each contained a single T-DNA insertion. Strain UOM DT5, however, carried a single T-DNA insertion as well as an additional partial insertion of 957 bp corresponding to the trp3 terminator located on a different contig. The sizes of the T-DNA insertions were 4,940 bp in UOM DT1 and 3,972 bp in UOM DT5, whereas the insertions in UOM STC1R2 and UOM STC4R2 measured 5,631 bp and 5,662 bp, respectively.

The specific genomic position of the T-DNA insertions in *D. tanaceti* UOM DT1, was identified downstream of gene G21 (Figure 3. A), which showed the highest similarity to a DNA repair and recombination protein rad5C from *Stagonosporopsis vannaccii* (NCBI accession: KAJ4988673.1; query coverage: 84%; per. identity: 68.91%). Alignment with the wild type revealed a 16 bp deletion at the insertion site (Figure 3. B). In *D. tanaceti* UOM DT5, the complete T-DNA was inserted within gene G25 (Figure 3. C), which showed highest similarity to a GMC oxidoreductase (GMCox) from *Stagonosporopsis vannaccii* (accession: KAJ4983148.1; query cover: 93%; per. identity: 77.61%). A comparison with the wild type identified a seven bp deletion and a 657 bp inversion downstream the T-DNA, and an 11 bp inversion and four bp deletion upstream the T-DNA, within GMCox (Figure 3. B). Additionally, a 957 bp region corresponding to the T-DNA terminator was located between genes G16 and G17 (Figure 3. C). BLAST results indicate that G16 shared the highest similarity with an uncharacterised protein (M421DRAFT_330253) from *Didymella exigua* (accession number: XP_033443501.1; query coverage: 49%; percentage identity: 82.45%,). On the other hand, G17 showed highest similarity to a hypothetical protein (E8E12_003031) from *Didymella heteroderae* (accession number: KAF3039328.1; query coverage: 82%; percentage identity: 53.11%)

**Figure 3.**
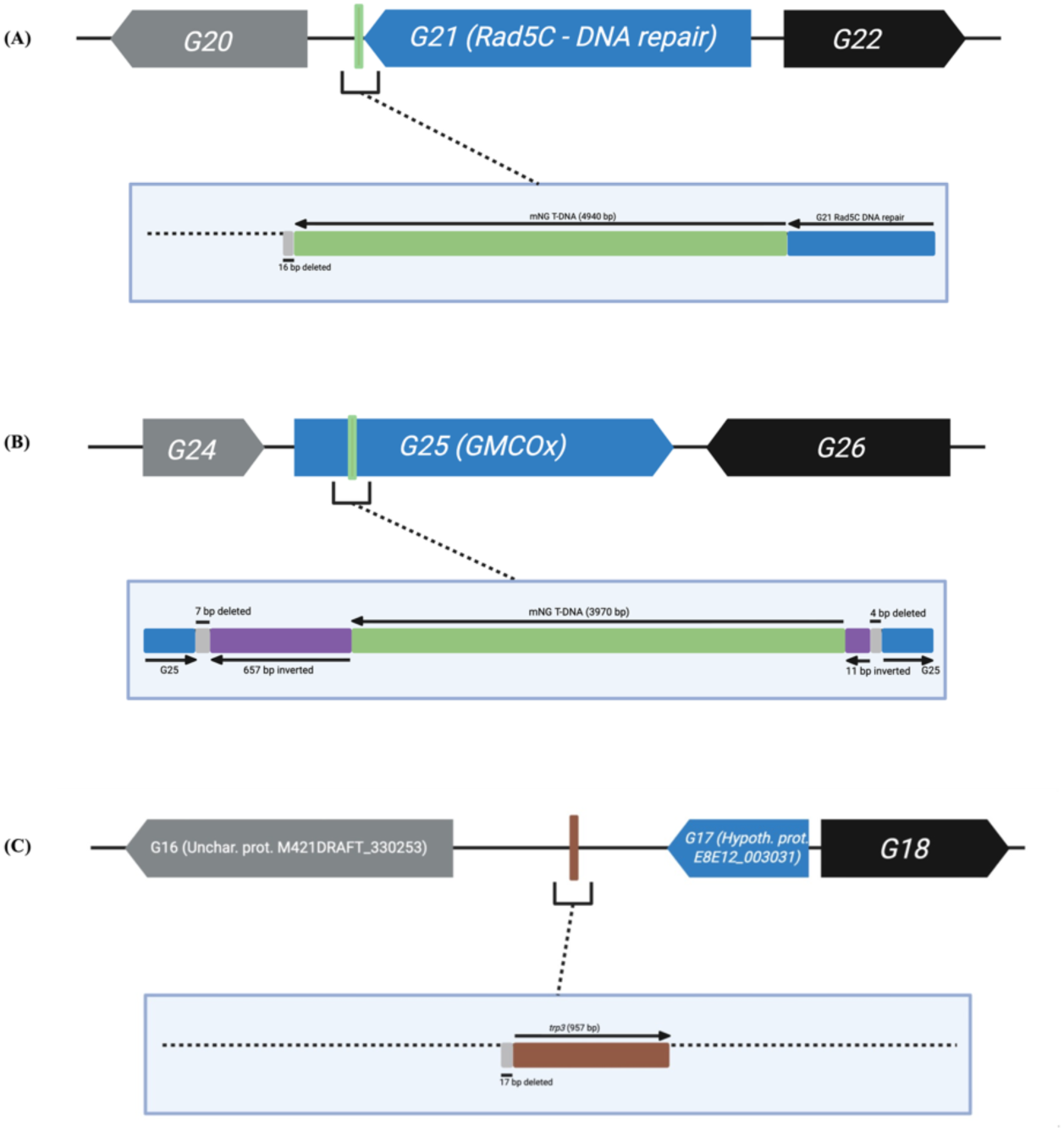
T-DNA insertion sites in *Didymella tanaceti* transformants. (A) Schematic representation of the T-DNA insertion site (green line) in strain UOM DT1. Alignment against the wild type showed a 16 bp deletion at the insertion site. (B) Schematic representation of the T-DNA insertion site (green line) in strain UOM DT5. Alignment with the wild type revealed an 11 bp deletion and a 657 bp inversion downstream of the insertion site, as well as an 11 bp inversion and a 4 bp deletion upstream of the insertion site. (C) Schematic representation of a partial T-DNA insertion in strain UOM DT5, comprising 957 bp of the trp3 terminator. Alignment against the wild type showed a 17 bp deletion at insertion site.

In *S. tanaceti* UOM STC1R2, the T-DNA was inserted within gene G31 (Figure 4. A), which showed the highest similarity to a Rheb small monomeric GTPase from *S. vannaccii* (NCBI accession: KAJ4987483.1; query coverage: 100%; per. identity: 75.32%). A 539 bp deletion was present at the insertion site (Figure 4. B). In *S. tanaceti* UOM STC4R2, the T-DNA was inserted downstream within gene G2 (Figure 4. C), which showed similarity to a hypothetical protein (SVAN01_07780) from *S. vannaccii* (accession: KAJ4986721.1; query coverage: 31%; per. identity: 88.73%). No deletions were detected at this insertion site (Figure 4. D).

**Figure 4.**
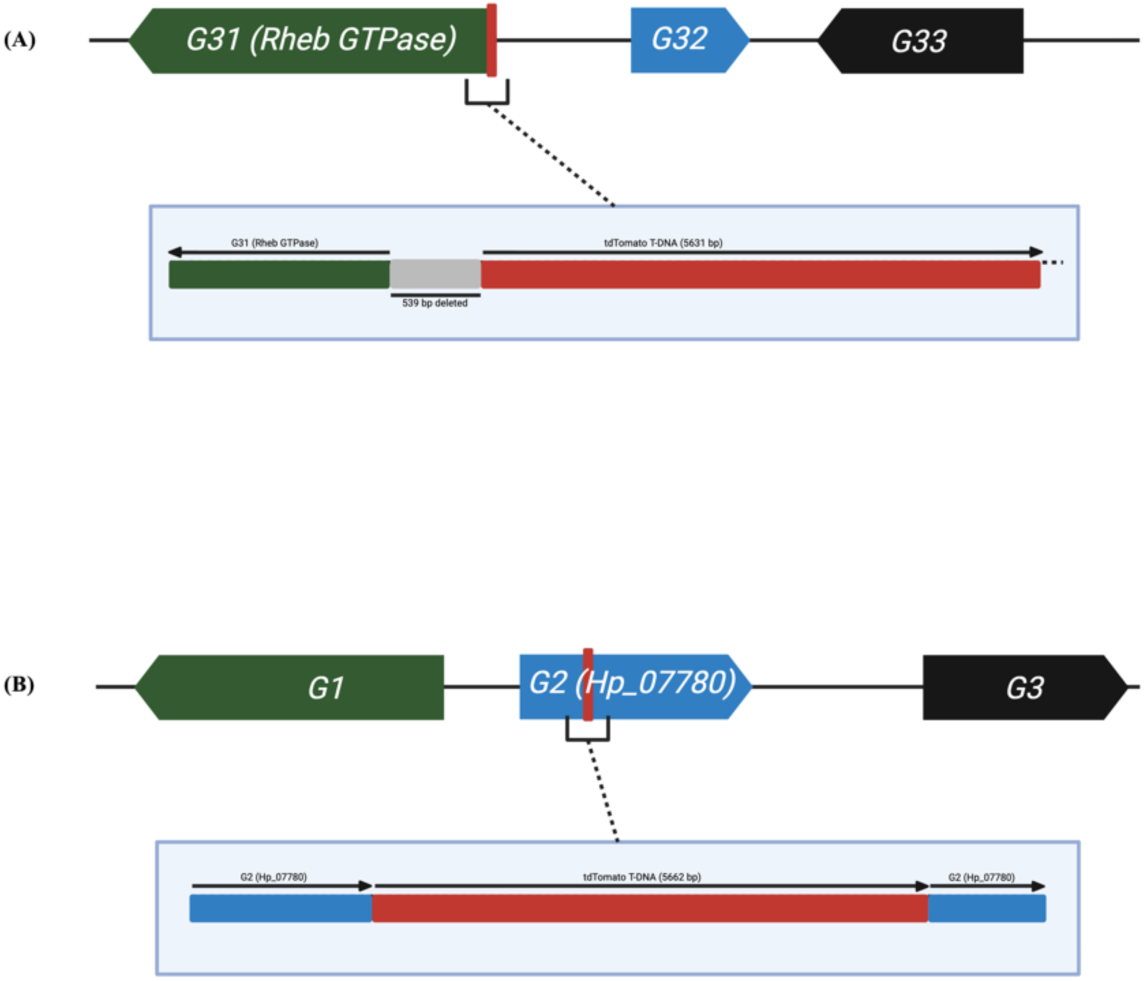
T-DNA insertion sites in *Stagonosporopsis tanaceti* transformants. (A) Schematic of the T-DNA insertion site (red line) in strain UOM STC1R2. Alignment with the wild type revealed a 539 bp deletion at the insertion site. (B) Schematic of the T-DNA insertion site (red line) in strain UOM STC4R2. Alignment with the wild type showed no deletions at the insertion site.

#### Wild type vs transformed strains

Initial symptoms were observed 5 DAI for both the *D. tanaceti* wild type strain and the transformed strain UOM DT5, appearing as dark-brown necrotic lesions on leaf surfaces and margins. Not all inoculated leaves developed visible symptoms. By 7 DAI, necrotic lesions on symptomatic leaves had enlarged and, in some cases, coalesced (Figure 5. A,B).

**Figure 5.**
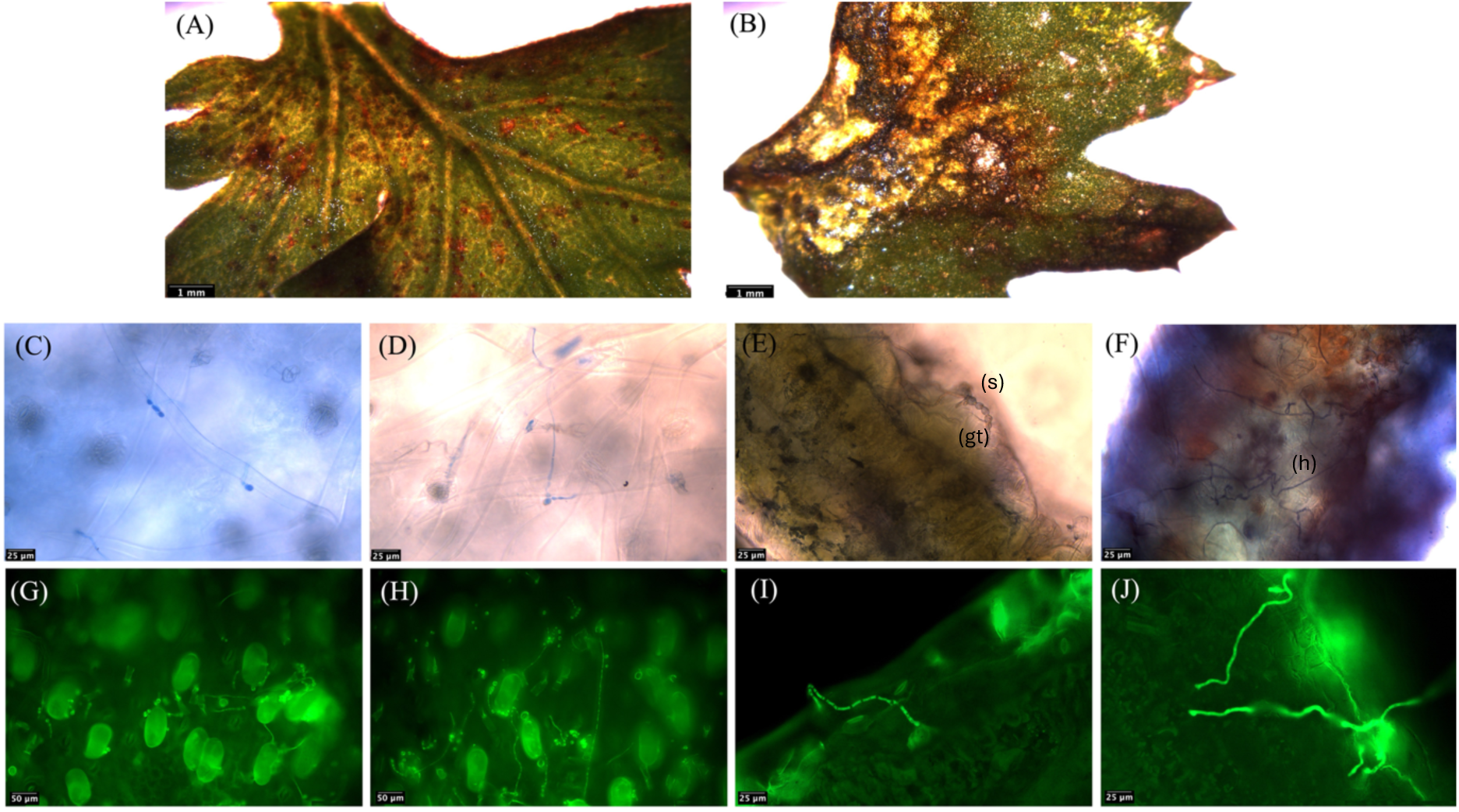
Tan spot lesions on pyrethrum leaves caused by *Didymella tanaceti* wild type and transformed strains, and subsequent disease progression. (A) Lesions caused by the non-transformed strain BRIP 61988 at 7 days after inoculation (DAI). (B) Lesions caused by the transformed strain DT5 at 7 DAI. (C-F) Disease progression of the wild type isolate at 24, 48, and 72 hours after inoculation (HAI), and 5 DAI, shown from left to right. The letter “s” indicates the spore, “gt” (E), and “h” indicates the fungal hyphae (F). (G-J) Disease progression of the transformed strain UOM DT5 at the same time points. Panels C, D, G, and H show the adaxial surface of infected leaves, while panels E, F, I, and J show transverse sections of infected leaves. Scale bars: A–B = 1 mm; C–J = 25 and 50 μm.

Spore germination was detected 24 hours after inoculation (HAI) in both wild type and transformed strain. In the wild type, spores typically formed swollen short chains before germ tube emergence (Figure 5. C), a feature also observed in the transformed strain UOM DT5 (Figure 5. G). At 48 HAI, both strains produced hyphae extending across the leaf surface, without evidence of specialized infection structures (Figure 5. D, H). By 72 HAI, penetration from the epidermis into the mesophyll was observed (Figure 5. E, I), and by 5 DAI, both strains had colonised the mesophyll through the middle lamella (Figure 5. F, J). No differences in germination pattern, penetration timing, or tissue colonisation were observed between wild type and transformed strains.

For *S. tanaceti*, initial symptoms appeared at 4 DAI as dark necrotic lesions at the margins. By 7 DAI, necrotic lesions extended across the entire or partial section of the leaf surface, irrespective of the strain (Figure 6. A, B). Symptomatic leaves were transversally sectioned at approximately 50-100 µm and observed under a compound microscope using the DsRed fluorescent filter at 200× and 400× magnification. The characteristic red fluorescence of UOM STC1R2 was detected in mycelium infecting the mesophyll at 3 DAI (Figure 6. A); however, sample thickness and tissue autofluorescence complicated visualisation.

**Figure 6.**
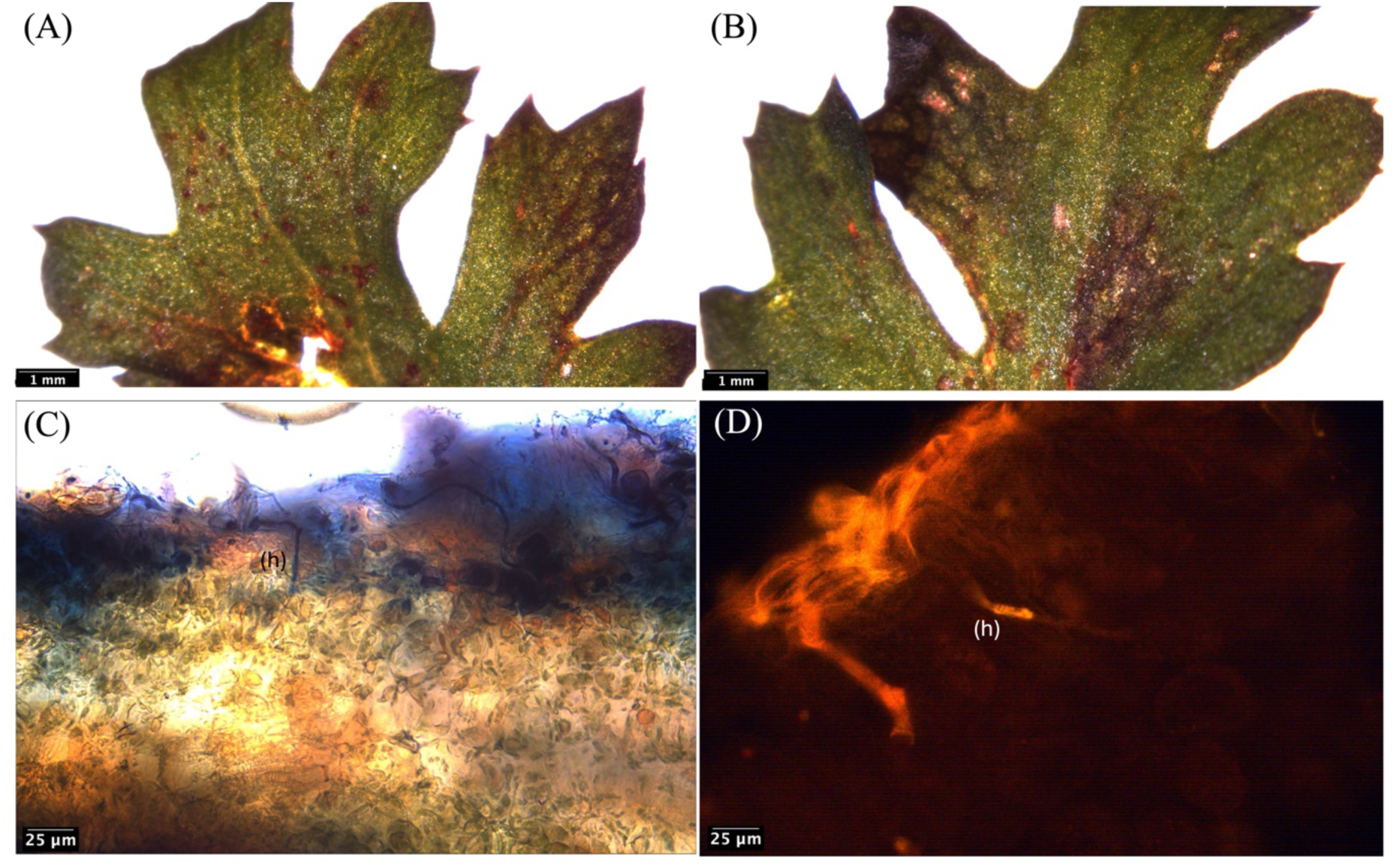
Lesions and tissue colonisation of pyrethrum leaves caused by *Stagonosporopsis tanaceti* wild type and transformed strains. (A) Lesions caused by the wild type strain at 7 days after inoculation (DAI). (B) Lesions caused by the transformed strain UOM STC1R2 at 7 DAI. (C) Transformed strain UOM STC1R2 and (D) wild type strain in transverse sections of infected leaves at 3 DAI. Letter “h” in C, D indicate fungal hyphae. Scale bars: A-B = 1 mm; C-D = 25 μm.

### Co-infection of transformed *D. tanaceti* and *S. tanaceti*

Co-inoculated leaflets were successfully cross-sectioned using a razor blade. Sections mounted in droplets of RO water were examined under a compound microscope equipped with DsRed and YFP filter sets. Fluorescence microscopy allowed clear differentiation of the two pathogens within the same tissue, with UOM STC1R2 (*S. tanaceti*) visualised via DsRed fluorescence and UOM DT5 (*D. tanaceti*) via YFP fluorescence. This approach demonstrated that both pathogens could be simultaneously identified and spatially distinguished in co-infected leaf tissues.

## Discussion

Understanding the infection process and disease cycle of a pathogen is fundamental for developing effective disease management strategies enabling targeted interventions during critical infection windows, thereby enhancing control efficacy (42). The infection biology of *Stagonosporopsis tanaceti* on pyrethrum is well characterised: the pathogen can originate from seedborne inoculum, causing symptoms in emerging seedlings or persisting as a latent infection within the host (11, 12). However, the disease cycle and infection process of *D. tanaceti* on pyrethrum remain largely unknown. Therefore, fluorescently labelled variants of *D. tanaceti* were generated to facilitate the investigation of its infection process, colonisation of the pyrethrum leaf mesophyll, and interactions with the other common pathogen of pyrethrum, *S. tanaceti*.

Fluorescently labelled strains of *D. tanaceti* and *S. tanaceti* were generated for the first time using plasmids pMAI31 and pMAI32, enabling visual tracking of both pathogens during the initial phase of infection. Most *Agrobacterium tumefaciens*-mediated transformation (ATMT) studies in filamentous fungi have focused on investigating gene function (21, 22, 43). However, the use of ATMT to insert fluorescent markers has also been applied, albeit to a lesser extent, to study pathogen infection dynamics and to enable visual tracking and differentiation of pathogens within host tissues (13, 15, 25, 44). Plasmids pMAI31 and pMAI32 were previously used to transform *Leptosphaeria maculans* successfully (45) demonstrating their utility for generating transgenic, fluorescently-labelled fungal strains.

Single-copy T-DNA insertions were obtained for both *D. tanaceti* and *S. tanaceti* under the ATMT conditions used. Whole-genome sequencing of *D. tanaceti* strains UOM DT1 and UOM DT5 and *S. tanaceti* strains UOM STC1R2 and UOM STC4R2 revealed that each transformant contained a single, T-DNA insertion. In addition to the T-DNA integration, strain UOM DT5 also carried a partial insertion consisting of 957 bp of the *L. maculans* trp3 terminator sequence located in a different genomic node. Although the T-DNA fragments identified in the transformed strains did not span the entire T-DNA contained between the right and left border of each plasmid, the sequences found in the transformed strains retained all the essential elements required for expression of the fluorescent markers. ATMT facilitates the random integration of T-DNA into the host genome and is often effective at producing single-copy insertions; however, multiple copies and tandem repeats can also occur (22, 46). In *Didymella rabiei*, a pathogen of chickpea from the family Didymellaceae, tandem repeat integrations have been reported (15), whereas in *Didymella lentis*, both single and multiple insertions have been documented (25).

Insertional mutagenesis resulted in T-DNA integration either adjacent to or within four distinct genes in the transformants. In strain UOM DT1, T-DNA was inserted immediately downstream of a Rad5-family gene (*rad5C*). The *rad5C* in UOM DT1 is a putative Rad5-like protein predicted to function in an error-free post-replication repair pathway, acting downstream of *Rad18* to mediate template switching and replication fork restart (47-49). Based on structural annotation, the downstream T-DNA insertion leaves the predicted *rad5C* open reading frame intact, consistent with the absence of detectable effects on growth or morphology. However, to accurately determine whether the insertion affects gene expression, RNA-seq or targeted genotoxic stress assays would be required.

In strain UOM DT5, the T-DNA was inserted within the coding sequence of a GMC oxidoreductase (*GMCox*) gene, as indicated by structural annotation. The GMC oxidoreductase family comprises a diverse group of enzymes, including both secreted and intracellular members with metabolic functions (50, 51). In plant-pathogenic fungi, several secreted oxidoreductases have been associated with lignocellulose degradation and host colonisation (51, 52). Potentially, the absence of any detectable effect on growth rate or virulence following disruption of this locus could reflect functional redundancy with other GMC-family homologues in the genome or indicate that the disrupted gene encodes an intracellular enzyme not directly involved in pathogenicity.

Inversions and deletions were detected on both sides of the T-DNA insertion site of strain UOM DT5. Unlike the rad5C locus in strain UOM DT1, the *GMCox* open reading frame in UOM DT5 is disrupted by the T-DNA, representing the most complex genomic rearrangement observed among the transformants. Small deletions at T-DNA integration sites have been previously reported in *Agrobacterium*-mediated transformations of plants and filamentous fungi. While large-scale inversions and deletions at T-DNA integration sites are frequently reported in plants and filamentous fungi (53, 54), small deletions and inversions also occur (55, 56). These minor rearrangements are less commonly documented, but may be more prevalent than currently appreciated (57).

In the *S. tanaceti* transformed strain UOM STC1R2, the T-DNA insertion is associated with a 539 bp deletion immediately upstream of the *RhebGTPase* gene (G31). The RhebGTPases are small GTP-binding proteins and key regulators of the TORC1 (Target of Rapamycin) signalling pathway, controlling processes like cell growth, nutrient sensing, and development in filamentous fungi (58, 59). The observed deletion may affect the start codon or promoter region, potentially altering transcription or translation efficiency. In strain UOM STC4R2, the T-DNA insertion occurs within a genomic region annotated as a hypothetical protein (*SVAN01_07780*) from *S. vannaccii*, without any deletions or inversions detected at the insertion site. Although the function of this region is unknown, this insertion illustrates that *Agrobacterium*-mediated transformation can target diverse genomic regions. The absence of structural rearrangements suggests that this insertion is a good example of a relatively simple integration event compared with the more complex rearrangements observed in the other transformants of *D. tanaceti* and *S. tanaceti*.

The fluorescence intensity values reported (Figures 3. A, B) closely reflected the patterns observed under fluorescence microscopy. Among the *D. tanaceti* transformed strains, fluorescence levels varied, with strain UOM DT5 exhibiting the highest intensity. Similar fluorescence intensity fluctuations in genome-integrated EGFP have been previously reported in *Cochliobolus heterostrophus* (60) and *Nigrospora* sp. (38). In contrast, *S. tanaceti* strains showed highly consistent fluorescence intensity across all strains, which were not statistically different. These results indicate that, despite some variation in intensity levels, the respective fluorescent proteins are stably expressed and functionally active in the transgenic lines of *D. tanaceti* and *S. tanaceti*, suggesting successful transformation and no apparent loss of reporter activity across strains. The greater variation observed in *D. tanaceti* could be attributed to differences in T-DNA integration sites and species-specific factors influencing promoter activity and protein stability.

The *in vitro* morphology and growth rate of the transformant remained similar to the wild type for most strains of both species, suggesting that T-DNA insertion did not disrupt genes essential for these traits in most cases. Growth rate was assessed immediately after confirming fluorescence, and strains exhibiting noticeably slower growth than the wild type were excluded from host interaction studies. Such differences in growth rate could indicate that the T-DNA insertion occurred within, or near, genes involved in growth regulation or related metabolic pathways (61, 62). Slower-growing strains were deemed unsuitable for further experiments because they would not accurately reflect the timing of the infection process and host-pathogen interaction *in planta*.

Direct epidermal penetration, rather than stomatal entry, was observed in *D. tanaceti* during infection of pyrethrum leaves. This infection process mirrors that of the wild type strain and confirms that transformation did not affect the pathogenicity of strain UOM DT5, as both showed comparable disease progression on detached leaves. Direct epidermal penetration has also been reported in *S. tanaceti* (41) and in several necrotrophic and hemi-biotrophic fungi, including *Botrytis cinerea* and *Colletotrichum* spp., where penetration pegs form beneath appressoria or directly from hyphae without stomatal involvement (63, 64). Although stomatal penetration was not observed in this study, further investigation is needed to determine whether *D. tanaceti* can also utilise this pathway, particularly since the closely related species *Didymella rabiei* is capable of both direct epidermal and stomatal entry (15).

Transverse sections of pyrethrum leaves infected with both the wild type and the transformed strain UOM STC1R2 of *S. tanaceti* revealed infection hyphae colonising parenchymatic tissue at 3 DAI. This indicates that the transformed strain progressed through the same infection stages as the wild type, consistent with previously reported timelines. Bhuiyan *et al.* (41) described the infection process of *S. tanaceti* in pyrethrum leaves using conventional histopathology methods, showing that germination occurred approximately 12 HAI and mesophyll invasion by around 54 HAI. The agreement between these findings and our observations suggests that transformation did not disrupt pathogenic development or tissue invasion dynamics in *S. tanaceti*, supporting UOM STC1R2 as a reliable model for fluorescence-based tracking of infection in pyrethrum. Overall, the comparable infection dynamics between wild type and transformed strains in both species confirm that the transformation process did not alter pathogenic development, validating the use of these fluorescent strains for *in planta* infection studies.

Pyrethrum leaves were successfully co-inoculated with *D. tanaceti* strain UOM DT5 and *S. tanaceti* strain UOM STC1R2, enabling simultaneous tracking of their colonisation patterns. By 7 DAI, both pathogens were observed invading parenchymatic tissue through the middle lamella, occupying similar spatial niches within the leaf. This spatial overlap in the mesophyll indicates the potential for direct competition for resources. However, no direct interaction between the two fungi was observed at this stage of infection. Experimental work in *Populus* demonstrated that competition for mesophyll cells between a *Melampsora* rust fungus and mites influenced pathogen antagonism by leaf endophytic fungi (65). The observation of both pathogens surrounding the same lesion (Figure 10 A) raises the possibility that symptom expression in co-infected leaves may result from synergic or antagonistic interactions rather than from the activity of a single pathogen. These findings highlight the value of fluorescently tagged strains as tools for investigating pathogen-pathogen interactions *in planta* and lay the groundwork for future studies aimed at quantifying competitive fitness and determining the ecological consequences of co-infection in pyrethrum.

The green, round structures observed in the fluorescent images in Figure 7. E, F correspond to glandular trichomes distributed across the surface of pyrethrum leaves, which can naturally fluoresce at multiple wavelengths (66-68). Their intrinsic fluorescence could present a challenge when visualising the pathogen on the leaf surface, particularly from the adaxial view, as trichome signals may overlap with pathogen-derived fluorescence. Plants naturally contain autofluorescent molecules such as chlorophyll, lignin, and flavonoids, which emit fluorescence primarily in the red (∼680-740 nm) and green (∼500-550 nm) regions of the spectrum (69). Moreover, infection can induce the accumulation of additional fluorescent compounds in leaves (70), which can potentially fluoresce under the microscope when appropriate excitation and emission conditions are used. Chlorophyll fluorescence itself is known to change in response to infection, reflecting physiological alterations in the plant (71). Furthermore, fluorescence has been associated with infection sites, likely due to the localised production of plant-derived compounds as part of the defence response (72). The natural autofluorescence spectra of healthy and diseased plant tissues largely overlap with the detection windows of the fluorescent reporter proteins employed in this study. However, the emission intensity of mNeonGreen (∼517 nm) was substantially higher than the green autofluorescence from leaf tissue, allowing clear differentiation between pathogen-derived fluorescence and background signals. Similarly, tdTomato (∼581 nm) generated a markedly stronger emission than the background red fluorescence from chlorophyll, which peaks at longer wavelengths. This contrast likely results from the strong, narrow emission peaks of the reporter proteins, in contrast to the broader and weaker autofluorescence characteristic of plant tissues.

**Figure 7.**
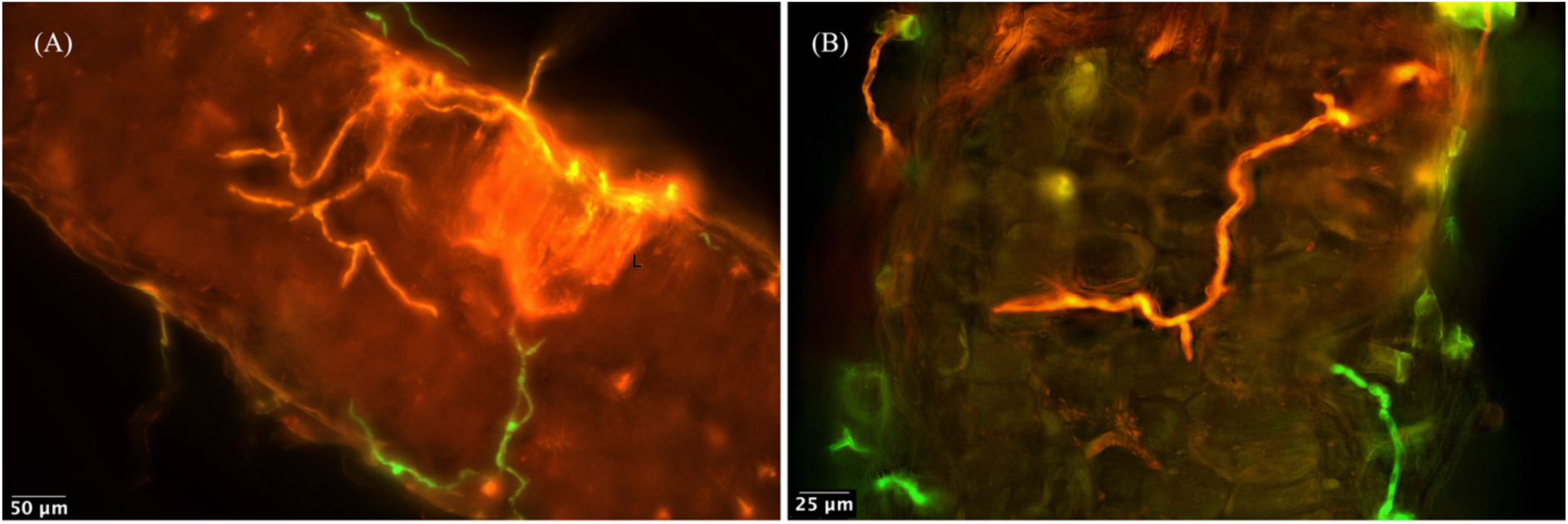
Cross-sections of pyrethrum leaves co-infected with *Stagonosporopsis tanaceti* strain UOM STC1R2 and *Didymella tanaceti* strain UOM DT5 at 200× magnification. Orange fluorescence corresponds to UOM STC1R2, and green fluorescence corresponds to UOM DT5. The letter “L” indicates a necrotic lesion (A).

The fluorescent *D. tanaceti* strains represents a powerful tool to investigate pathogen entry points and colonisation patterns by inoculating different plant tissues and tracking infection. This approach could also reveal whether the pathogen moves systemically, as demonstrated for *Colletotrichum graminicola*, which spreads from roots to aerial tissues (16). Similarly, GFP labelling of *A. lentis* enabled detailed characterisation of *ascochyta* blight progression in lentil (15). Together with the present findings, these studies illustrate the value of fluorescent protein reporters for elucidating infection biology in poorly characterised pathogens. Moreover, using different reporter genes like *tdTomato* and *mNeonGreen* could enable visualisation of co-infection events, providing a more accurate reflection of the complex pathogen interactions that occur under natural field conditions.

## Acknowledgements

This research was supported by the University of Melbourne Scholarship and Botanical Resources Australia Pty Ltd. Thanks to Camilla Langlands-Perry for her support during the transformation processes.

## Supplemental figures

**Figure 1.**
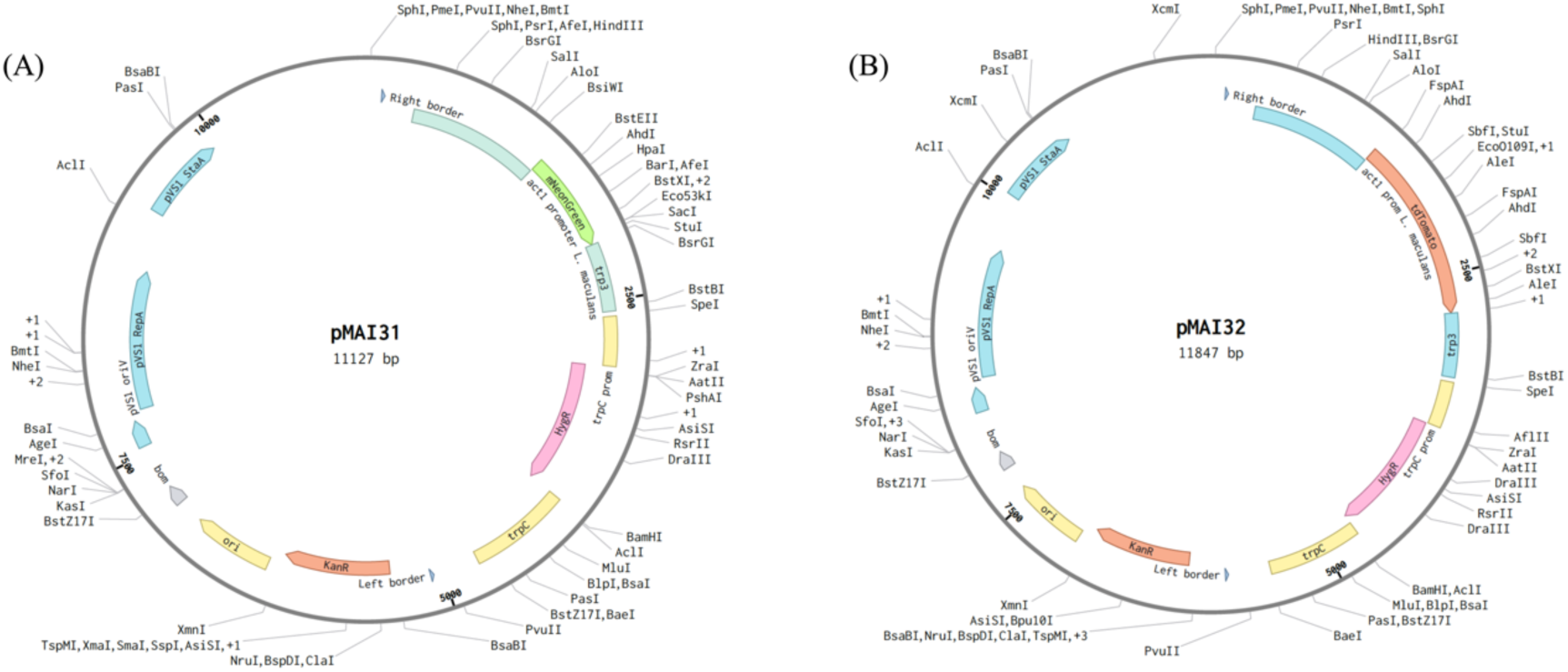
Schematic representation of plasmids (A) pMAI31 carrying the mNeonGreen fluorescent protein gene and (B) pMAI32 carrying the tdTomato fluorescent protein gene used to transform *Didymella tanaceti* and *Stagonosporopsis tanaceti*, respectively.

**Figure 2.**
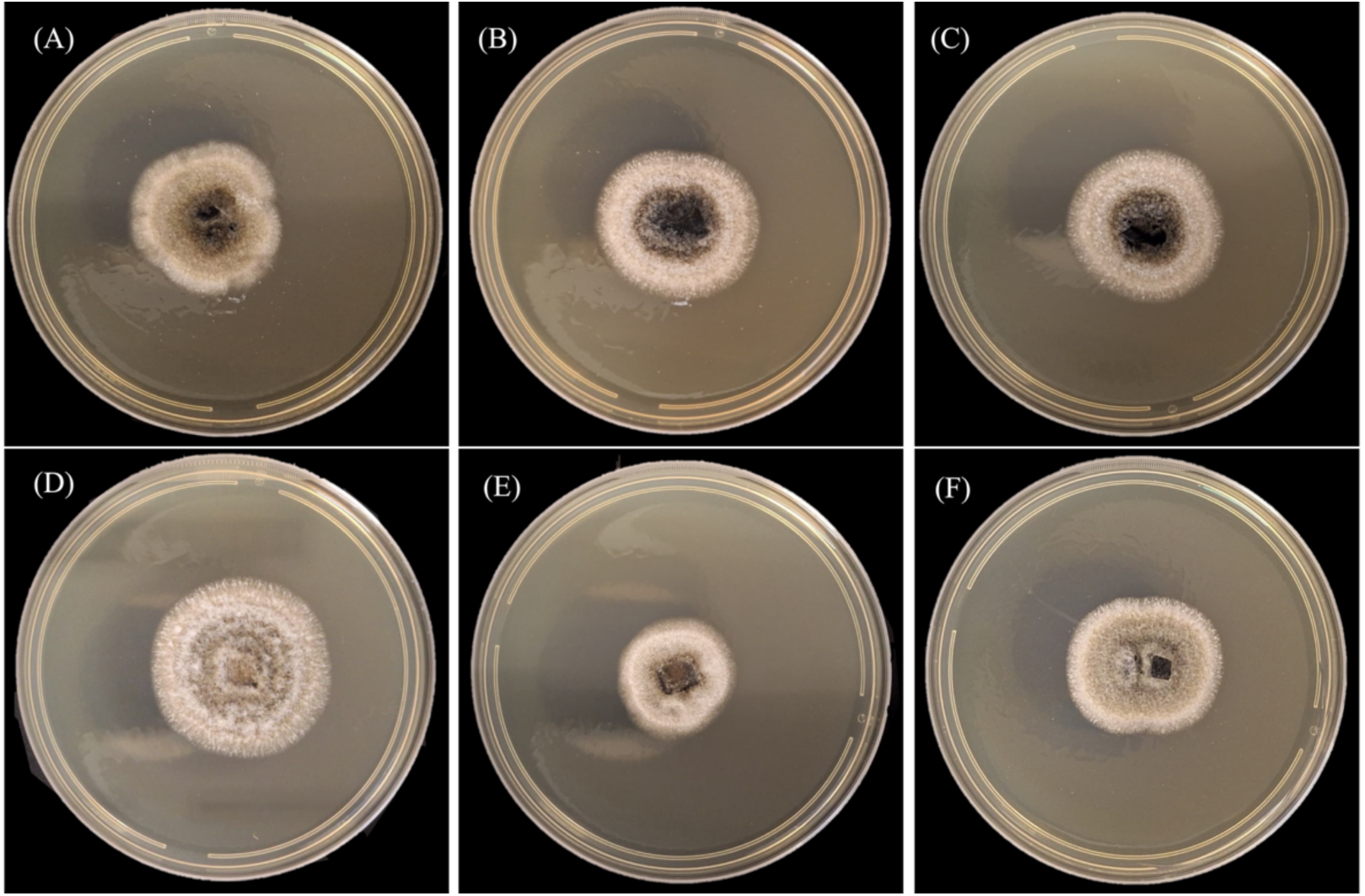
Morphological characterisation of wild type and selected transformed strains of *Didymella tanaceti after six days on potato dextrose agar. (A) Wild type strain BRIP 61988; (B-F) Transformed strains UOM DT1, UOM DT2, UOM DT3, UOM DT4 and UOM DT5*.

**Figure 3.**
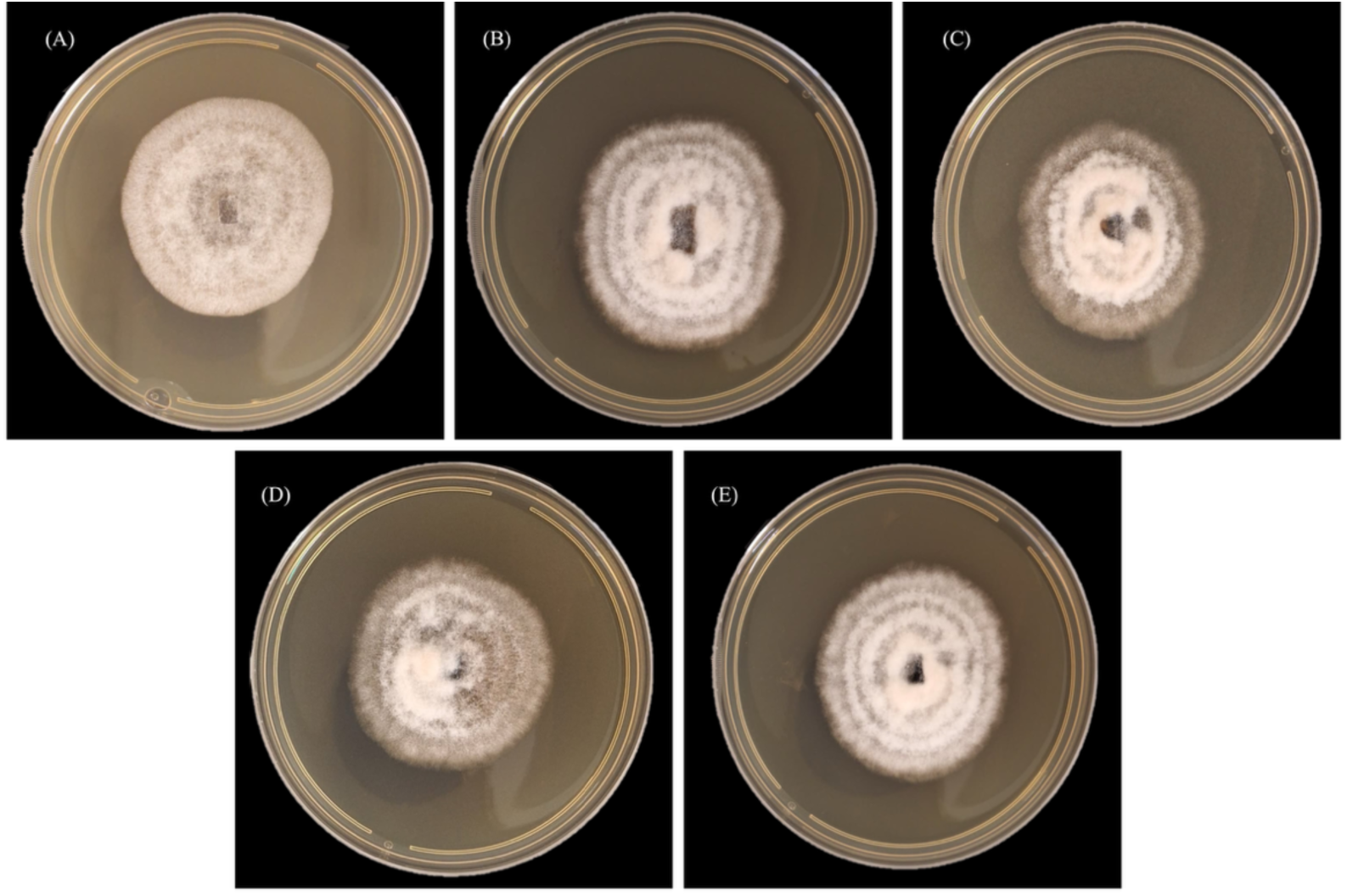
Morphological characterisation of wild type and selected transformed strains of *Stagonosporopsis tanaceti after six days on potato dextrose agar. (A) Wild type strain UOM ST2; (B-E) Transformed strains UOM STC1R2, UOM STC2R2, UOM STC3R2 and UOM STC4R2*

## Supplementary tables

**Table 1.** Growth rates (mean ± SD, mm) of *Stagonosporopsis tanaceti* wild type (UOM ST2) and transformed strains (UOM STC1R–STC4R2) measured at Day 3, 6, and 9. Letters indicate statistically significant differences between strains within each time point based on Tukey’s post-hoc test (p < 0.05)

| *Day* | *Strain* | *mean_sd* |
| --- | --- | --- |
| *Day3* | UOM ST2 | 11.7 ± 0.6 b |
|  | UOM STC1R2 | 10.7 ± 0.6 ab |
|  | UOM STC2R2 | 10.3 ± 1.2 a |
|  | UOM STC3R2 | 10.7 ± 0.6 ab |
|  | UOM STC4R2 | 10.7 ± 0.6 ab |
| *Day6* | UOM ST2 | 24.7 ± 0.6 a |
|  | UOM STC1R2 | 23.7 ± 0.6 a |
|  | UOM STC2R2 | 24.7 ± 0.6 a |
|  | UOM STC3R2 | 24.7 ± 0.6 a |
|  | UOM STC4R2 | 24 ± 0 a |
| *Day9* | UOM ST2 | 35 ± 0 a |
|  | UOM STC1R2 | 35 ± 0 a |
|  | UOM STC2R2 | 35.3 ± 0.6 a |
|  | UOM STC3R2 | 35 ± 0 a |
|  | UOM STC4R2 | 34.7 ± 0.6 a |

**Table 2.**
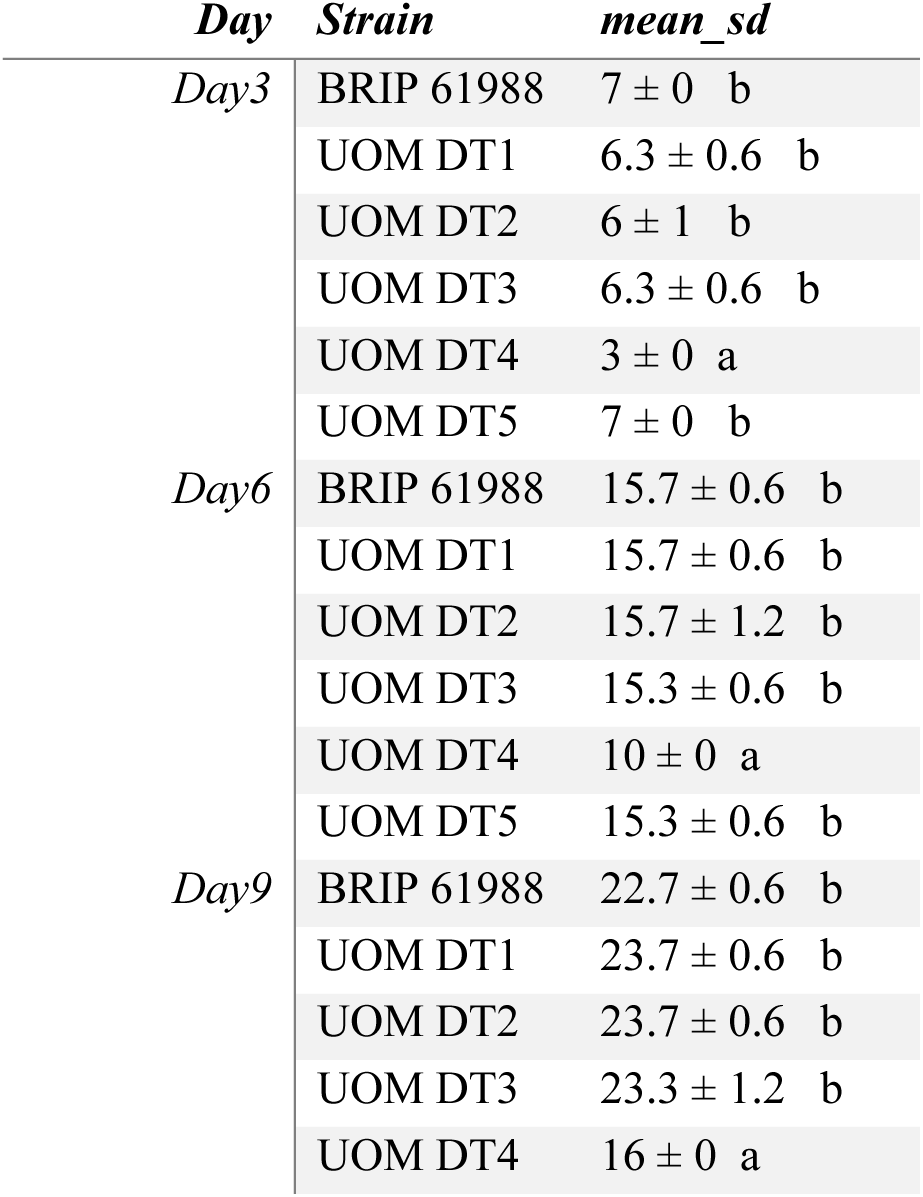
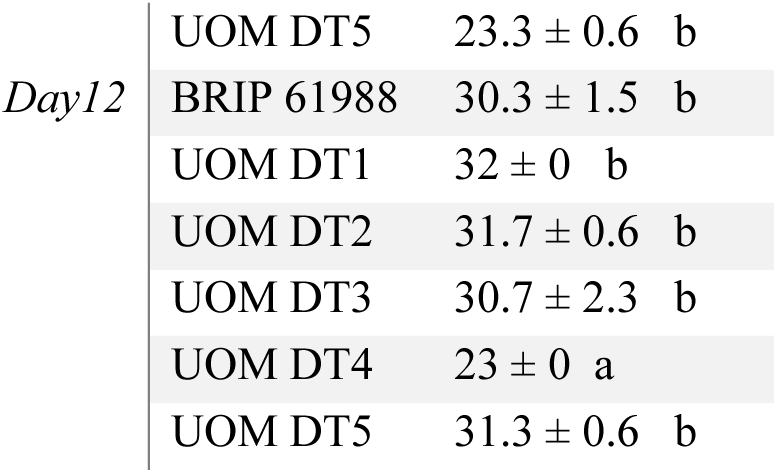
Growth rates (mean ± SD, mm) of *Didymella tanaceti* wild type (BRIP 61988) and transformed strains (UOM DT1–DT5) measured at Day 3, 6, 9, and 12. Letters indicate statistically significant differences between strains within each time point based on Tukey’s post-hoc test (p < 0.05).

