## Supplementary material for "Elucidating pathogen interactions in *Tanacetum cinerariifolium* (pyrethrum) using fluorescently labelled *Didymella tanaceti* and *Stagonosporopsis tanaceti*": Suplemental materials

### Supplementary materials

#### Potato dextrose agar (PDA), water agar (WA) and V8 media preparation

**PDA** was prepared by weighing 37 g of commercial PDA powder (Difco™) and dissolving it in 1 L of RO water. The mixture was stirred until fully homogenised and then autoclaved at 121 °C for 20 minutes.

WA was prepared by adding 20 g of agar powder (DifcoTM) to 1 L of RO water. Upon mixing, the media was autoclaved at 121 °C for 20 minutes. Similarly, Potato Dextrose Agar (PDA) media was prepared by weighing 37 g of commercial PDA powder (DifcoTM) and adding it to 1 L of RO water. The mixture was shaken until uniformity and autoclaved for 20 minutes at 121 °C.

V8 agar (Bhuiyan et al., 2017) was prepared by adding 200 ml of V8 vegetable juice filtered with a cheesecloth to 800 ml of RO water. Drops of 10 M NaOH were added to the mixture until a pH of 6.25 was achieved. Then, 20 g of agar were added, shaken until uniform and autoclaved at 121°C for 20 minutes.

#### ATMT induction media, antibiotics and overlay media preparation

<u>Cefotaxime, hygromycin and acetosyringone preparation.</u>

A 100 mg/mL cefotaxime stock solution was prepared, filter-sterilized, and stored at 4 °C. Hygromycin was used as a pre-made 50 mg/mL stock solution throughout the experiment. Acetosyringone was prepared as a 10 mM stock by dissolving 0.0196 g in 500 µL of dimethyl sulfoxide (DMSO) (Sigma Aldrich, USA). Short-term storage was at 4 °C, and long-term storage was at −21 °C.

Table 1. MM salts for induction media (IM)

| Formula | Chemical Name | Amount per litre (g) |
|---|---|---|
| $KH_2PO_4$ | Potassium dihydrogen phosphate | 3.625 |
| $K_2HPO_4$ | Dipotassium hydrogen phosphate | 5.125 |
| NaCl | Sodium chloride | 0.375 |
| $MgSO_4 \cdot 7H_2O$ | Magnesium sulfate heptahydrate | 1.250 |
| $CaCl_2 \cdot 2H_2O$ | Calcium chloride dihydrate | 0.165 |
| $FeSO_4 \cdot 7H_2O$ | Ferrous sulfate heptahydrate | 0.0062 |
| $(NH_4)_2SO_4$ | Ammonium sulfate | 1.250 |

*Note: each salt was entirely dissolved before adding the next one*

Preparation of 1M MES for IM

1 M MES buffer (pH 5.3) was prepared to use in Induction Media (IM) by dissolving 7.7 g of MES (2-(N-morpholino) ethanesulfonic acid) in approximately 45 mL of distilled water. The pH was adjusted to 5.3 using 5 M KOH, and the solution was brought to a final volume of 50 mL with distilled water. The buffer was filter sterilised and added directly to molten IM once the medium had cooled to ~50 °C.

IM media preparation

To prepare 1 L of IM agar, 400 mL of 2.5× MM salts, 0.6 g of glucose, 5 mL of glycerol, 545 mL of distilled water, and 20 g of agar were mixed. The media was autoclaved at 121°C for 20 minutes. Once cooled to 50 °C, 50 mL of sterile 1 M MES stock (pH 5.3) and 500 µL of acetosyringone (AS) were added. After mixing, the media was poured Petri plates and stored at 4 °C as required.

##### Overlay media preparation

The overlay medium for transformation, was V8 media for *S. tanaceti* and PDA for *D. tanaceti*. After autoclaving, the media was cooled to approximately 50 °C. Then, cefotaxime and hygromycin were added to achieve final concentrations of 100 µg/mL and 50 µg/mL, respectively. This was done by adding 200 µg/mL cefotaxime stock solution and 100 µg/mL hygromycin stock solution at a 1:1,000 dilution into the molten medium. The medium was mixed gently but thoroughly before being poured into Petri plates.
